# Cholesterol differentially regulates α-synuclein binding across membrane packing regimes

**DOI:** 10.64898/2026.05.21.726870

**Authors:** Orianna H. Kou, Brian H. Kim, David H. Johnson, Wade F. Zeno

**Author notes:** **Corresponding Author:** Wade F. Zeno –.

## Abstract

α-Synuclein (αSyn) is an intrinsically disordered protein that preferentially binds anionic membranes with lipid packing defects. Cholesterol is an abundant membrane component that regulates packing and organization within membranes, yet its effect on αSyn binding remains unclear as prior studies report both cholesterol-mediated enhancement and suppression. Here, we investigated whether these conflicting effects reflect differences in the intrinsic packing state of the phospholipid bilayer. Using a quantitative fluorescence microscopy-based binding assay, we measured αSyn binding preferences among reconstituted phosphatidylcholine/phosphatidylserine membranes with varied cholesterol content, lipid tail chemistry, and vesicle curvature. We found that cholesterol’s effect depended on the underlying packing regime of the membrane. In defect-rich membranes, cholesterol reduced αSyn binding, consistent with cholesterol tightening lipid packing and reducing αSyn-accessible defects. In membranes with intermediate defect content, cholesterol enhanced binding, whereas tightly packed membranes remained largely insensitive to cholesterol except when high cholesterol content was combined with high membrane curvature. Curvature further shaped these responses, with high curvature compressing cholesterol-dependent differences between membrane compositions. These results show that cholesterol does not universally promote or inhibit αSyn binding. Instead, cholesterol regulates αSyn-membrane interactions through a packing-regime-dependent mechanism shaped by both lipid tail chemistry and membrane curvature. This framework helps reconcile opposing reports in the literature and highlights membrane physical state as a key determinant of how cholesterol modulates αSyn binding.

**Significance:** αSyn is a membrane-binding protein associated with Parkinson’s disease, but the role of cholesterol in regulating its membrane interactions has remained unclear. Some studies report that cholesterol enhances αSyn association with membranes, whereas others show that cholesterol suppresses it. This work helps explain why both outcomes can occur. We show that cholesterol’s effect depends on the membrane’s underlying packing state: cholesterol can reduce, enhance, or have little effect on αSyn binding depending on the membrane environment. These findings shift the question from whether cholesterol is generally pro- or anti-binding to how cholesterol reshapes the membrane physical states that control αSyn association.

## Introduction

αSyn is a small, 140-residue, intrinsically disordered protein localized predominantly at neuronal synaptic terminals^1^, where its pathological aggregates are closely associated with synucleopathies such as Parkinson’s disease and Lewy body dementia^2–5^. Because αSyn aggregation is closely linked to its membrane interactions^3, 6–8^, a detailed understanding of the physicochemical determinants governing αSyn-membrane association is essential for understanding its pathological behavior.

Upon membrane binding, the N-terminus of αSyn transitions from a disordered conformation to an amphipathic α-helix, embedding its hydrophobic face into the bilayer while its positively charged residues engage anionic phospholipids electrostatically^9, 10^ (Fig. 1A). Since binding is facilitated by membrane insertion, αSyn has particularly high affinity for membranes that are highly curved^11^ and rich in lipid packing defects^12–20^, which are local regions of exposed lipid tails that arise from lipid splaying or non-ideal packing. Cholesterol, an abundant component of biological membranes^21^, alters bilayer order and packing^22^ and can therefore influence αSyn-membrane interactions^23, 24^. However, the relationship between cholesterol and αSyn binding remains unresolved^25, 26^. Some studies report positive correlations between cholesterol levels and αSyn membrane association^27–30^, whereas others show that cholesterol suppresses αSyn binding^31–33^. Additional studies suggest that cholesterol effects are context-dependent, varying with membrane composition, phase behavior, and the type of model membrane used^34, 35^. In some cases, cholesterol-dependent binding was non-monotonic^36^, further indicating that cholesterol effects cannot be captured by a simple increase-or-decrease model. This complexity extends to αSyn aggregation, with some studies showing that cholesterol enhances aggregation propensity^35, 37–39^, while others suggest that cholesterol can reduce aggregation under specific conditions^36^. Together, these findings highlight the need for a physical framework that explains when cholesterol promotes, suppresses, or fails to alter αSyn-membrane interactions.

**Figure 1:**
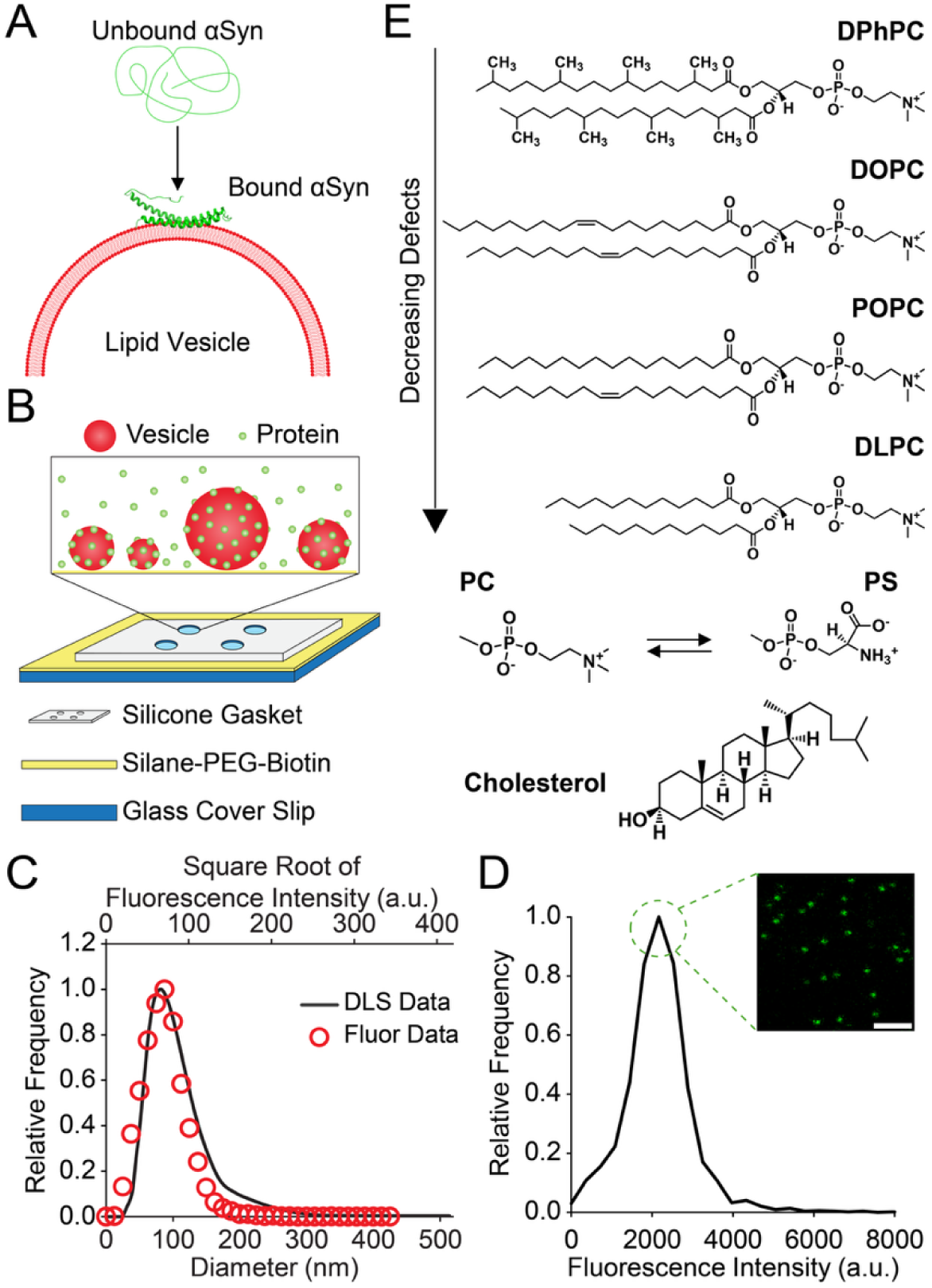
A tethered-vesicle assay to monitor αSyn membrane binding as a function of lipid composition. (A) Schematic of αSyn binding to a membrane surface. The schematic is not drawn to scale and is not intended to reflect αSyn’s exact membrane insertion depth. (B) Experimental configuration of the tethered-vesicle assay. (C) Representative vesicle diameter calibration using lipid fluorescence intensity and DLS measurements. (D) Representative single-molecule calibration to determine the fluorescence intensity of αSyn monomers. Scale bar represents 2 μm. (E) Structures of the lipid species used in our vesicle binding experiments.

This complexity likely arises because cholesterol can alter membrane packing in multiple, context-dependent ways. Cholesterol often orders phospholipid acyl chains and reduces membrane fluidity, which can limit the packing defects needed for αSyn insertion^34, 40, 41^. However, cholesterol can also disrupt overly tight lipid packing and alter local membrane organization^42^, meaning its effect may depend on the initial packing state of the bilayer. Therefore, cholesterol’s impact on αSyn binding may not depend simply on cholesterol content, but also on the intrinsic defect landscape of the membrane. This possibility remains poorly explored, particularly in systems that titrate cholesterol while holding membrane charge constant across tail-chemistry-defined packing regimes.

Here, we used a quantitative fluorescence microscopy assay to investigate αSyn membrane binding as a function of cholesterol content and membrane packing defects. To isolate these contributions, we used small unilamellar vesicles (SUVs) composed of phosphatidylcholine (PC) and phosphatidylserine (PS) at a fixed PC:PS molar ratio of 3:1 while varying cholesterol content across lipid tail chemistries that span high- and low-defect regimes. We found that cholesterol suppresses αSyn binding in high-defect membranes, enhances binding in intermediate-defect membranes, and has little effect in low-defect membranes, except when high cholesterol is combined with high membrane curvature. These findings establish a packing-regime framework in which cholesterol’s regulation of αSyn binding depends on the physical state of the membrane.

## Materials and Methods

### Materials

Sodium phosphate monobasic and sodium phosphate dibasic were from Sigma-Aldrich. ATTO 647N-labeled 1,2-Dipalmitoyl-sn-glycero-3-phosphoethanolamine (DPPE) and ATTO 488-maleimide were from ATTO-TEC. 1,2-diphytanoyl-sn-glycero-3-phosphocholine (DPhPC), 1,2-diphytanoyl-sn-glycero-3-phospho-L-serine (sodium salt) (DPhPS), 1,2-dioleoyl-sn-glycero-3-phosphocholine (DOPC), 1,2-dioleoyl-sn-glycero-3-phospho-L-serine (sodium salt) (DOPS), 1-palmitoyl-2-oleoyl-glycero-3-phosphocholine (POPC), 1-palmitoyl-2-oleoyl-sn-glycero-3-phospho-L-serine (sodium salt) (POPS), 1,2-dilauroyl-sn-glycero-3-phosphocholine (DLPC), 1,2-dilauroyl-sn-glycero-3-phospho-L-serine (sodium salt) (DLPS), and cholesterol (plant) were from Avanti Research. Dipalmitoyl-decaethylene glycol-biotin (DP-EG15-biotin) was provided by D. Sasaki from Sandia National Laboratories, Livermore, CA^43^. Biotin-PEG-Silane (5 kDa) and mPEG-Silane (5 kDa) were purchased from Laysan Bio, Inc. Silicone gaskets were purchased from Grace Bio-Labs.

### α-Synuclein Expression, Purification, and Fluorescent Labeling

Fluorescent αSyn was prepared exactly as described previously^15, 44^. In brief, a BL21 (DE3) *E. coli* co-expression system was used to obtain N-terminally acetylated αSyn with a Y136C mutation. This post-translational modification was included as it represents the physiological form of the protein^45–47^. αSyn was then purified using ammonium sulfate precipitation, anion-exchange chromatography, and size-exclusion chromatography. Finally, αSyn was concentrated and fluorescently labeled at the cystine residue using ATTO 488-maleimide. UV-vis spectroscopy was used to confirm a 1:1 dye:protein labeling efficiency.

### Tethered-Vesicle Assay

SUVs were prepared exactly as described previously^15, 44^. SUV formulations in this work were prepared as 100 μM (lipid concentration) stock solutions that contained PC:PS lipids at a 3:1 molar ratio, 0-40 mol% cholesterol, 0.5 mol% DP-EG15-biotin, and 0.5 mol% DPPE-ATTO 647N.

Sample imaging wells were prepared as described previously^15, 44, 48, 49^. Briefly, imaging wells were created by adhering silicone gaskets with 5 mm-diameter holes to glass coverslips, onto which biotinylated PEG-silane was covalently attached (Fig. 1B). SUVs containing DP-EG15-biotin were then tethered to the surface using NeutrAvidin. After 10 minutes of incubation, excess unbound SUVs were rinsed away, and αSyn was then incubated with the tethered vesicles.

Imaging was performed on a Leica Stellaris 5 confocal microscope with a Leica HC PL APO 63x oil-immersion objective (NA 1.4). ATTO 488-labeled αSyn was excited at 488 nm (emission 493-558 nm); ATTO 647N-labeled SUVs were excited at 638 nm (emission 643-749 nm). All scans used a 400 Hz line frequency with optical zoom set to yield 70 nm x 70 nm pixels. All samples were imaged in buffer containing 20 mM sodium phosphate and 150 mM NaCl (pH 7.0).

### Image Calibration and Analysis

Image calibration and analysis followed procedures described previously^15, 44^. Fluorescent puncta were identified in the lipid channel using cmeAnalysis^50^ and retained only if present across three consecutive imaging frames. Vesicle diameters were estimated by calibrating lipid-channel fluorescence against DLS measurements (i.e., lipid fluorescence ∝ vesicle area ∝ diameter^2^) (Fig. 1C). αSyn binding was quantified from colocalized protein-channel fluorescence using single-molecule ATTO 488 calibrations adjusted for detector gain (Fig. 1D).

### Mean Binding Response Calculation

The mean binding response was calculated as the concentration-averaged number of bound proteins over the tested αSyn concentration range (equation 1).

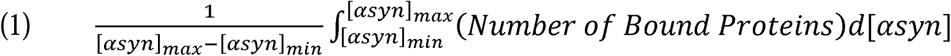

Specifically, the area under each binding curve was divided by the concentration interval, yielding a single binding metric with units of number of bound proteins.

The integral was approximated numerically using the trapezoidal rule.

## Results

### Defect-dependent membrane binding of α-synuclein

To determine how lipid tail chemistry regulates αSyn membrane binding, we prepared vesicles composed of PC and PS lipids at a fixed 3:1 molar ratio while varying the lipid tails (Fig. 1E). Four lipid tail chemistries were selected to span a range of expected membrane packing defect densities: dilauroyl (DL), 1-palmitoyl-2-oleoyl (PO), dioleoyl (DO), and diphytanoyl (DPh), with DPh membranes expected to be the most defect-rich and DL membranes expected to be the most tightly packed. This ordering is supported by prior experimental^51^ and simulation^14, 15, 52^ studies showing that these lipid tail features produce systematic differences in packing defect density. αSyn binding was quantified using a tethered vesicle assay (Figs. 1B-D, see methods). This technique enabled simultaneous measurement of vesicle size and αSyn binding for individual vesicles. Representative fluorescence micrographs showed a clear dependence of αSyn binding on lipid tail chemistry (Fig. 2A). Protein-channel intensity was highest for DPhPC:DPhPS vesicles and decreased progressively through DOPC:DOPS, POPC:POPS, and DLPC:DLPS vesicles. The resulting distributions of bound αSyn per vesicle followed the same trend observed in the images, with the highest binding to DPh membranes and the lowest binding to DL membranes (Fig. 2B). Across lipid compositions, individual vesicle diameters spanned approximately 30-150 nm, while the population mean diameter for each condition fell between approximately 80 nm and 100 nm.

**Figure 2:**
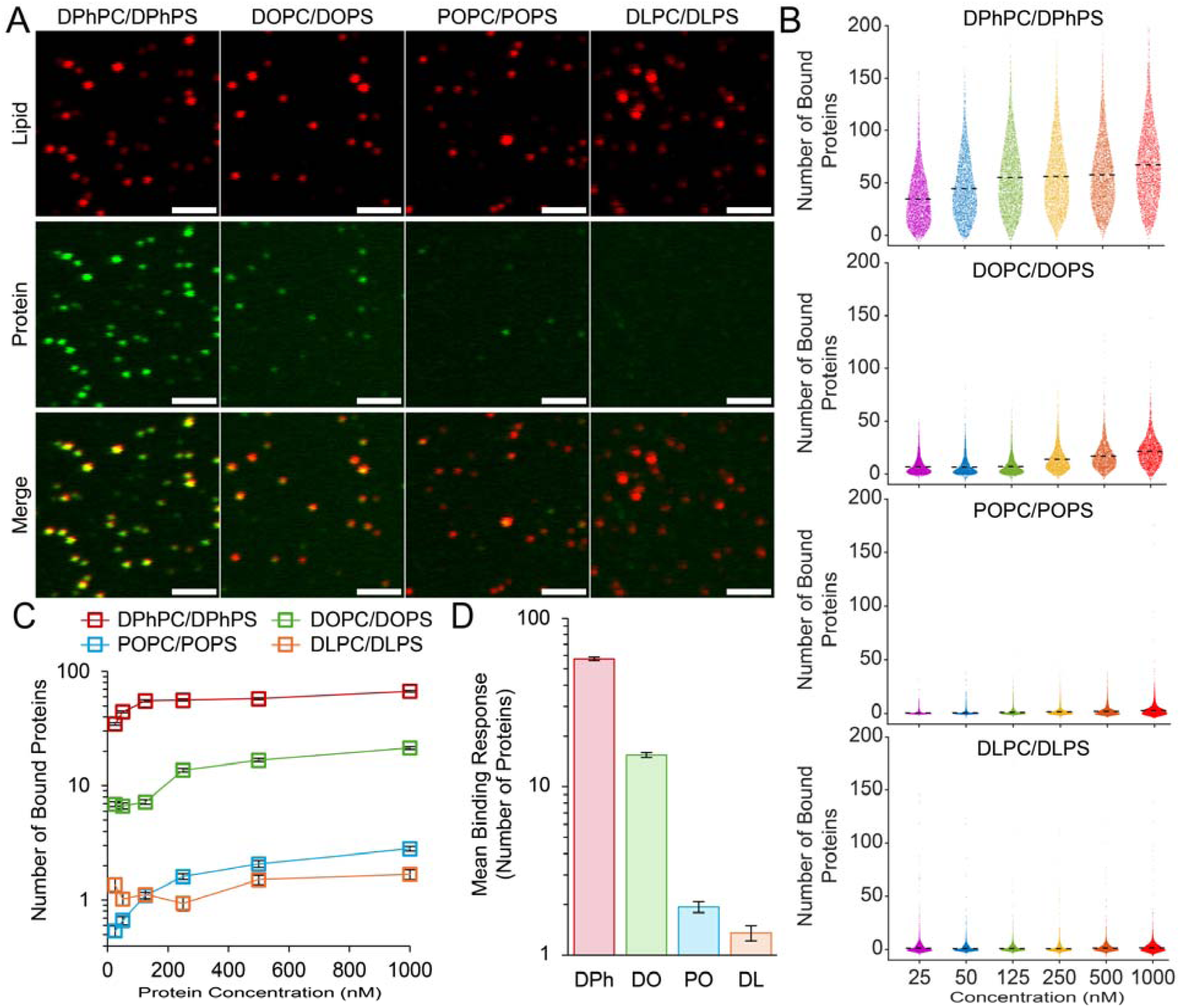
Lipid packing defects enhance αSyn binding to membranes. (A) Representative fluorescence micrographs of 1 μM αSyn bound to lipid vesicles. Scale bars represent 2 μm. (B) Distribution of bound proteins on the entire vesicle population spanning 30-150 nm diameter. Dashed lines represent the mean values of the distributions. A small number of outlier vesicles were visually omitted from the plots because they fell outside the selected y-axis range. Specifically, 10 DPhPC/DPhPS vesicles and 1 DOPC/DOPS vesicle across all concentration ranges. (C) Overlaid binding curves for αSyn at various lipid compositions. N = 2936-6663 vesicles for each point on the binding curve. Error bars represent the 99% confidence interval of the population averages in B. (D) Mean binding response of αSyn determined from the binding curves in C. Error bars represent the propagated error from C.

Binding curves were generated from the average number of bound proteins per vesicle (Fig. 2C). The mean binding responses showed a consistent rank order across protein concentrations: DPh > DO > PO > DL (Fig. 2D). These results establish the expected defect-dependent binding hierarchy in our assay and provide the baseline membrane packing regimes used to evaluate cholesterol-dependent changes in αSyn binding.

### Cholesterol differentially modulates α-synuclein binding

We next examined how cholesterol affects αSyn binding across membranes with different inherent packing defect regimes. For each composition, cholesterol was added while maintaining a fixed 3:1 PC:PS molar ratio, allowing us to vary cholesterol content without changing the overall anionic lipid fraction. DO, PO, and DL membranes were tested from 0 to 40 mol% cholesterol, whereas DPh membranes were limited to 25 mol% cholesterol because higher cholesterol fractions produced insoluble lipid mixtures and unreliable vesicle preparations.

Cholesterol produced distinct binding responses across the four baseline packing regimes (Fig. 3). In the highest-binding DPh membranes, increasing cholesterol reduced αSyn binding (Fig. 3A). In DO membranes, which showed intermediate binding in the absence of cholesterol, cholesterol increased αSyn binding, with the strongest enhancement at the highest cholesterol condition (Fig. 3B). PO membranes, which showed low baseline binding in the absence of cholesterol, also exhibited enhanced αSyn binding with added cholesterol, although this effect was weaker than in DO-based membranes (Fig. 3C). In the lowest-binding DL membranes, αSyn binding showed no clear cholesterol-dependent trend (Fig. 3D). Together, these results show that cholesterol does not uniformly enhance or suppress αSyn binding but instead produces distinct responses across membrane packing regimes.

**Figure 3:**
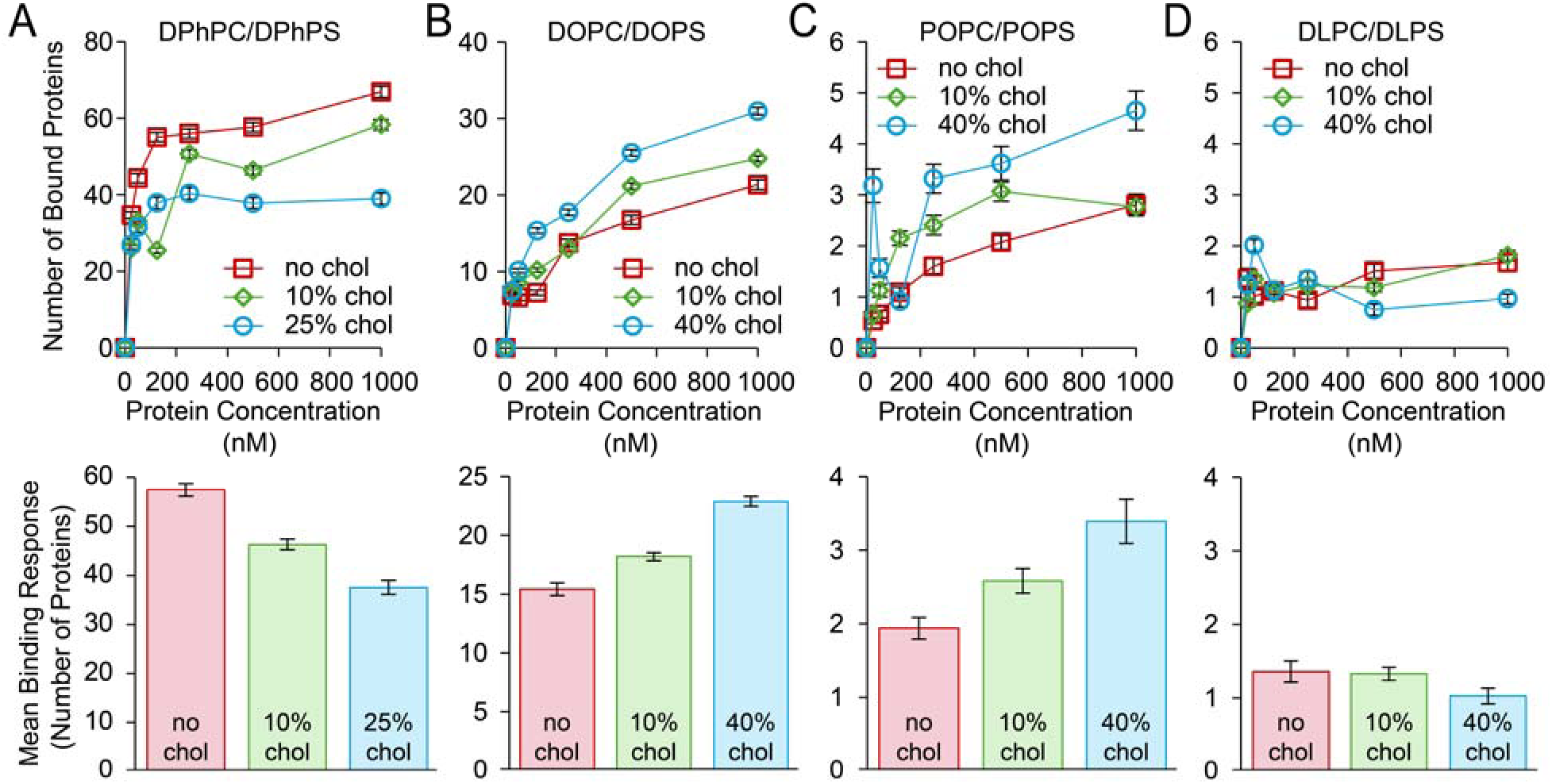
Cholesterol differentially modulates αSyn binding across membrane defect regimes. αSyn binding curves (top) and corresponding mean binding responses (bottom) in vesicles composed primarily of (A) DPh lipids (N = 1702-5292 vesicles per data point), (B) DO lipids (N = 2936-10495 vesicles per data point), (C) PO lipids (N = 1741-5813 vesicles per data point), or (D) DL lipids (N = 4685-6663 vesicles per data point). Error bars in binding curves represent the 99% confidence interval of the mean. Error bars in mean binding responses represent the propagated error from the binding curves.

### Impact of membrane curvature on cholesterol-dependent αSyn binding

To simultaneously compare cholesterol responses across membrane packing regimes and vesicle sizes, we plotted αSyn binding at 1 μM across the full 30-150 nm vesicle diameter range (Fig. 4A). This analysis reinforced the same composition-dependent responses observed in Figure 3: cholesterol decreased αSyn binding to DPh membranes, increased binding to DO membranes, and had little effect on DL membranes. For both DPh and DO membranes, the separation between cholesterol conditions increased at larger vesicle diameters (i.e., lower curvature), indicating that cholesterol-dependent effects were more pronounced at lower membrane curvature. Interestingly, PO membranes showed a distinct local binding enhancement near 50 nm diameter that diminished at larger diameters, suggesting that cholesterol enhances αSyn binding within this composition only in the high-curvature regime.

**Figure 4:**
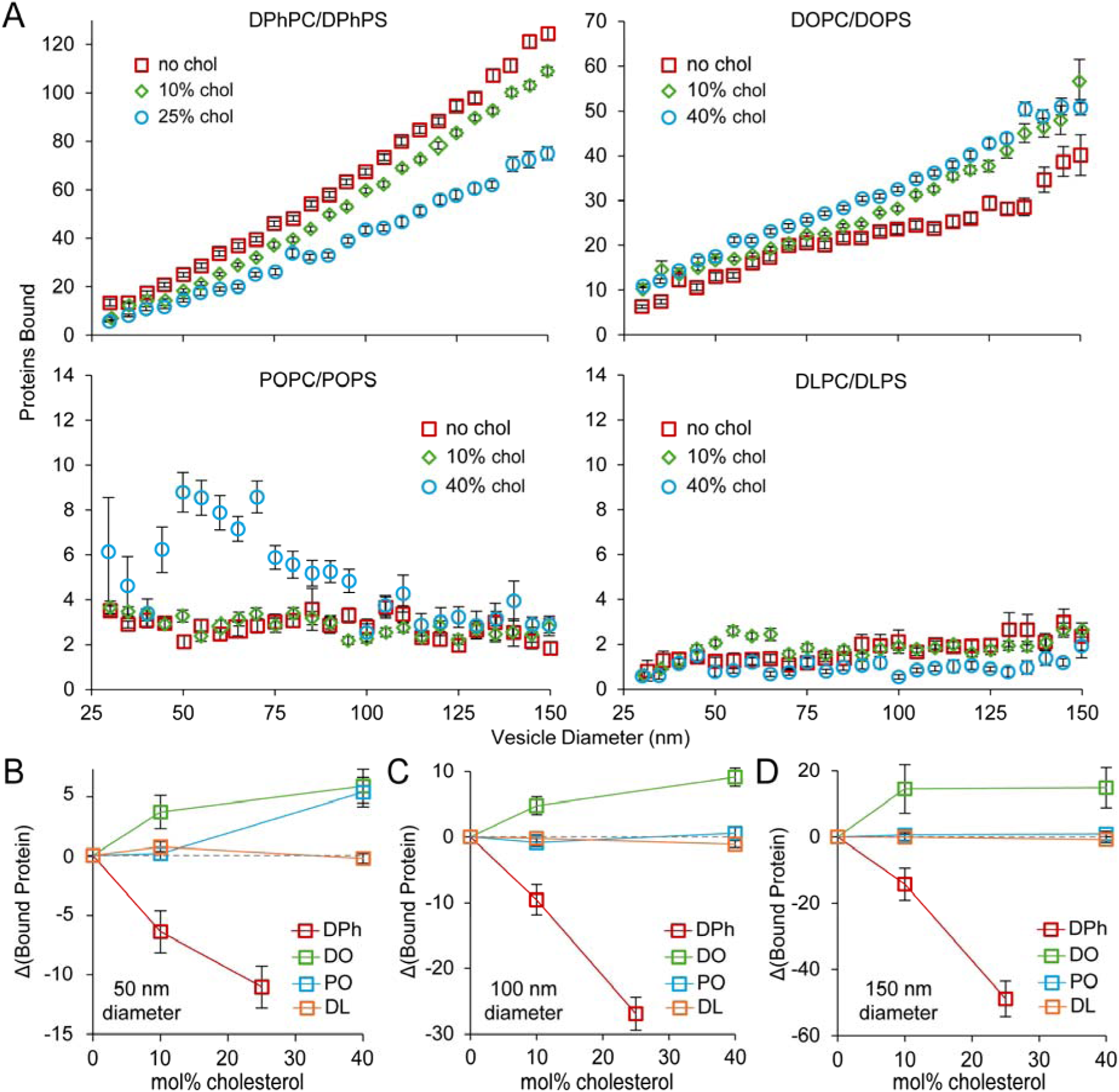
Membrane curvature dependence of cholesterol-regulated αSyn binding. (A) αSyn binding as a function of both vesicle diameter and cholesterol mole percentage. Data points represent 5-nm wide diameter bins centered every 5 nm from 30 to 150 nm (N= 13-974 vesicles per bin). Error bars represent the standard error of the mean. (B-D) Changes in the number of bound αSyn monomers on vesicles with average diameters of 50 nm (N = 142-1224 vesicles per bin), 100 nm (N = 315-2699 vesicles per bin), or 150 nm (N = 110-733 vesicles per bin). Bins in B-D are 10-nm wide and centered at 50, 100, and 150 nm. Error bars in B-D represent 99% confidence intervals propagated from corresponding bins for cholesterol-containing and cholesterol-free conditions. For all conditions in A-D, the bulk αSyn concentration was 1 μM.

We next summarized these trends by calculating the change in αSyn binding relative to the corresponding 0 mol% cholesterol condition for three distinct vesicle diameter populations (Figs. 4B-D). Cholesterol reduced αSyn binding to DPh membranes and enhanced binding to DO membranes across all vesicle sizes. In contrast, PO membranes showed enhanced binding only at 40 mol% cholesterol in 50 nm vesicles, while DL membranes remained largely insensitive to cholesterol. Together, these results show that membrane curvature modulates cholesterol-dependent αSyn binding, with the strongest curvature effect observed in PO membranes.

## Discussion

Our findings support a regime-dependent model for how cholesterol regulates αSyn binding to membranes. Rather than acting as a universal enhancer or suppressor, cholesterol produced distinct effects depending on the inherent packing defect state of the membrane. This dependence helps reconcile a literature in which cholesterol has been reported to promote αSyn membrane association, aggregation, or pathology in some contexts, while suppressing monomeric αSyn binding in others.

The decrease in αSyn binding observed in DPh membranes is consistent with the common interpretation that cholesterol can condense membranes and reduce the availability of packing defects. Several reconstituted studies have reported cholesterol-dependent decreases in αSyn binding, often attributed to increased lipid order, reduced fluidity, or reduced accessibility of hydrophobic defects^31–35^. Our DPh results are consistent with this model in a high-defect regime: when αSyn binding is already strong, adding cholesterol likely reduces the number or accessibility of productive insertion sites. The larger cholesterol-dependent separation observed at lower curvatures suggests that high curvature partially buffers this suppression by adding curvature-induced packing stress or defects. Thus, cholesterol suppresses binding in DPh membranes overall, but this effect is less pronounced when high curvature maintains additional αSyn-accessible defects.

The increased binding observed in DO membranes shows that cholesterol cannot be described only as a defect-plugging molecule. Instead, cholesterol enhanced αSyn binding in this intermediate-defect regime, suggesting that DO membranes occupy a packing state where cholesterol shifts the bilayer toward a more favorable binding environment. This finding aligns more closely with studies suggesting that cholesterol can enhance membrane binding, either monotonically^27, 29, 30^ or non-monotonically^36^. It may also help explain why raft-like or phase-separated cholesterol-containing membranes can support αSyn association even when cholesterol suppresses binding in simpler bilayers^25, 26, 34^. In these cases, cholesterol may create or stabilize membrane states that favor αSyn binding through local heterogeneity, altered headgroup spacing, domain boundaries, or sterol-associated contacts.

The PO and DL results define the boundaries of this cholesterol response. PO membranes showed enhancement only under high-cholesterol and high-curvature conditions, whereas DL membranes remained essentially nonresponsive. These findings argue against a simple model in which αSyn binds cholesterol-rich membranes because of cholesterol alone. If direct cholesterol recognition were sufficient, cholesterol would be expected to rescue binding in the weakest binding membranes. Instead, cholesterol-dependent enhancement appears to require a membrane that is already near a permissive packing state, or one that can be pushed toward that state by curvature.

This interpretation explains the parallel curvature dependence observed for DPh and DO membranes: high curvature compresses cholesterol-dependent differences by moving membranes toward a more defect-rich state. In DPh membranes, this curvature contribution partially offsets cholesterol-dependent suppression, whereas in DO membranes, it partially masks cholesterol-dependent enhancement because the highly curved membrane is already closer to a binding-permissive regime. At lower curvature, the influence of cholesterol becomes more apparent, causing the cholesterol conditions to separate more strongly. This interpretation is consistent with the broader defect framework in which electrostatics can support membrane recruitment, but productive αSyn insertion requires accessible packing defects^15, 16^.

This framework also helps separate direct membrane-binding effects from downstream cellular and aggregation phenotypes. Cellular studies often associate cholesterol-rich environments with increased αSyn membrane localization, accumulation, secretion, or pathology^27, 37^. Other studies suggest that cholesterol can influence αSyn aggregation or toxicity without clearly demonstrating that cholesterol directly enhances monomeric binding to a bilayer^35, 38, 39^. Our results suggest that such observations need not conflict with studies showing cholesterol-dependent binding suppression. Cholesterol may decrease monomeric binding in highly defect-rich membranes, enhance binding in intermediate regimes, or affect aggregation and propagation through mechanisms beyond simple membrane affinity, including changes in bound-state conformation, vesicle clustering, trafficking, lysosomal function, or lipid droplet biology^30, 32, 53^.

A limitation of the present study is that we do not directly measure how cholesterol changes membrane packing defects, lipid order, or lateral organization in each composition. Therefore, we cannot yet assign the DO enhancement to a specific physical mechanism. Cholesterol may increase the availability of αSyn-accessible defects in this regime, alter the size or lifetime of existing defects, promote membrane heterogeneity, or introduce weak sterol-specific interactions. Future simulations or measurements of lipid order and defect distributions across this cholesterol titration series will be needed to distinguish among these possibilities.

Overall, these results shift the cholesterol question away from whether cholesterol increases or decreases αSyn binding and toward identifying the membrane regimes in which each outcome occurs. Cholesterol suppresses αSyn binding in high-defect membranes (i.e. DPh lipids), enhances binding in intermediate-defect membranes (i.e., DO lipids), and has little effect in tightly packed membranes (i.e., DO or DL lipids), but can promote binding in PO membranes when high cholesterol is combined with high curvature. This regime-dependent model provides a simple physical framework for interpreting conflicting cholesterol effects in the αSyn literature and emphasizes that cholesterol regulates αSyn-membrane interactions through the membrane packing environment in which it is embedded. Importantly, this packing environment is shaped by both lipid tail chemistry and membrane curvature, which can introduce additional defects that alter the cholesterol response.

## Data availability

Data sets generated during the current study are available from the corresponding author upon reasonable request.

## Acknowledgments

Thank you to Jennifer C. Lee for providing the *pRK172 human* α*Syn(Y136C)* plasmid. Thank you to Peter J. Chung for providing the pNatB plasmid used to create N-terminally acetylated αSyn. This research was supported by the National Institutes of Health through R35GM147333 to O.H.K., D.H.J., and W.F.Z.

## Author contributions

O.H.K. and W.F.Z. conceived the project. O.H.K., B.H.K., and D.H.J. performed the experiments. O.H.K. and B.H.K. performed the analysis. O.H.K. and W.F.Z. wrote the manuscript with input from all authors.

